# Sparsity of population activity in the hippocampus is task-invariant across the trisynaptic circuit and dorsoventral axis

**DOI:** 10.1101/2024.09.17.613549

**Authors:** J. Quinn Lee, Matt Nielsen, Rebecca McHugh, Erik Morgan, Nancy S. Hong, Robert J. Sutherland, Robert J. McDonald

## Abstract

Evidence from neurophysiological and genetic studies demonstrates that *activity sparsity* – the proportion of neurons that are active at a given time in a population – systematically varies across the canonical trisynaptic circuit of the hippocampus. Recent work has also shown that sparsity varies across the hippocampal dorsoventral (long) axis, wherein activity is sparser in ventral than dorsal regions. While the hippocampus has a critical role in long-term memory (LTM), whether sparsity across the trisynaptic circuit and hippocampal long axis is task-dependent or invariant remains unknown. Importantly, representational sparsity has significant implications for neural computation and theoretical models of learning and memory within and beyond the hippocampus. Here we used functional molecular imaging to quantify sparsity in the rat hippocampus during performance of the Morris water task (MWT) and contextual fear discrimination (CFD) – two popular and distinct assays of LTM. We found that activity sparsity is highly reliable across memory tasks, wherein activity increases sequentially across the trisynaptic circuit (DG < CA3 < CA1) and decreases across the long axis (ventral < dorsal). These results have important implications for models of hippocampal function and suggest that activity sparsity is a preserved property in the hippocampal system across cognitive settings.

## Main text

Diverse evidence from genetic, anatomical, and physiological studies has revealed that *activity sparsity*, the proportion of active neurons in a population, varies significantly and systematically throughout the hippocampus (Chawla et al., 2005; Chawla et al., 2018; Jung & McNaughton, 1993; Thome et al., 2017; Witharana et al., 2016). Namely, the classical trisynaptic circuit (dentate gyrus (DG) → CA3 → CA1) is known to have the sparsest activity in the DG, and increasing levels of activity in CA3 and CA1, respectively (Chawla et al., 2005; Thome et al., 2017). Recent findings have also shown that sparsity varies across the dorsoventral (long) axis of the hippocampus (Chawla et al., 2018; Lee et al., 2019; Lee et al., 2022). These findings have important implications for theoretical models of neural computation in the hippocampus, wherein sparsity is thought to have a central role in memory capacity and fidelity (Cayco-Gajic & Silver, 2019; Marr, 1971). Despite the known importance of the hippocampus for long-term memory (LTM) (Lee et al., 2016; Teyler & DiScenna, 1985) and the role of sparsity in neural computation (Hunter et al., 2022; Thome et al., 2017), no studies to our knowledge have measured sparsity across the trisynaptic circuit and long axis of the hippocampus during performance of a memory task.

To address this gap in knowledge, we used functional molecular imaging to quantify sparsity across the trisynaptic circuit and long axis of the rat hippocampus during performance of two popular memory tasks. Namely, we trained rats to navigate to a hidden goal location in the Morris water task (MWT), or to avoid a fear-associated context in a contextual fear discrimination (CFD) paradigm (Figure 1; Detailed Methods) (Lee et al., 2022). Following performance of each task, we measured activity sparsity as the proportion of cells expressing *Arc* – a molecular marker of neural activity associated with learning and memory (Chawla et al., 2005; Guzowski et al., 1999; Lee et al., 2022). In doing so, we aimed to address if the pattern of activity sparsity in the hippocampus is task-dependent or invariant.

**Figure 1.**
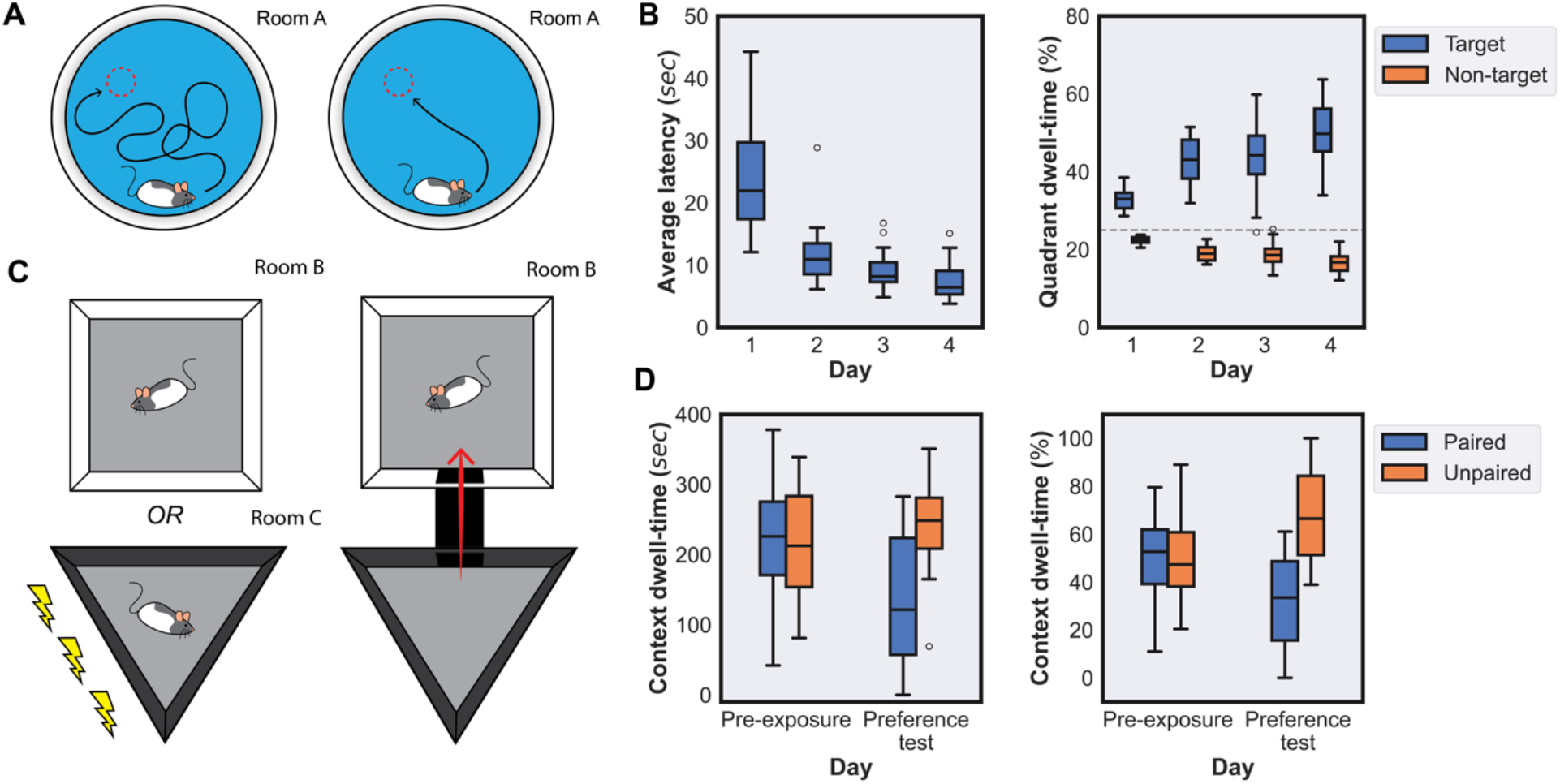
Expression of spatial and context fear memory in two tasks. **(A)** The schematic illustrates training in the MWT, wherein animals were trained over 4 days to locate a hidden platform from random start locations in a pool. **(B)** The boxplots show the average latency across of the entire behavioral cohort to reach the hidden platform on each day (left), and the percentage of dwell time in target versus non-target quadrants (right). We observed a significant eCect of day on average latency to reach the hidden platform (linear regression: *R*=-0.6886, *P*<0.0001), and a significant eCect of day (ANOVA: *F*(Day)=4.0457, *P*(Day)=0.0096), quadrant (ANOVA: *P*(Quadrant)<0.0001, *F*(Day X Quadrant)=16.1827), and day X quadrant interaction (ANOVA: *F*(Day X Quadrant)=16.1827, *P*(Day X Quadrant)<0.0001) in percentage of target versus non-target quadrant dwell time. Post-hoc related samples t-tests on each day of quadrant dwell time showed significant differences on Day 1 (*t*(1,11)=9.4850, *P*<0.0001), Day 2 (*t*(1,11)=9.7234, *P*<0.0001), Day 3 (*t*(1,11)=6.2742, *P*<0.0001), and Day 4 (*t*(1,11)=9.3653, *P*<0.0001). **(C)** The graphic illustrates training in and testing in the CFD task, wherein animals were first pre-exposed to two contexts, then conditioned with three foot-shocks on a single day to avoid one of two contexts upon subsequent context preference testing. **(D)** The boxplots show dwell time (left) and percentage of total dwell time (right) in each context during pre-exposure and preference test behavioral sessions. A two-way ANOVA revealed a significant eCect of day (*F*=4.5850, *P*=0.0349), context (*F*=10.1882, *P*=0.0019), and day X context interaction (*F*=13.7978, *P*=0.0003) on dwell time, and a significant eCect of context (*F*=21.7019, *P*<0.0001) and day X context interaction (*F*=26.2057, *P*<0.0001) for percent dwell time, but no eCect of day (*F*<0.0000, *P*=1.0000). Post-hoc related samples t-tests showed a significant difference in dwell time during the preference test (*t*=-4.3036, *P*=0.0003) but not pre-exposure (*t*=0.2803, *P*=0.7817), and a significant difference in percent dwell time during the preference test (*t*=-4.5130, *P*=0.0002) but not during pre-exposure (*t*=0.2533, *P*=0.8023).

To evaluate spatial learning in the MWT, we measured average latency to the goal location and the percentage of dwell time in the target versus non-target pool quadrants (Figure 1A-B). This revealed a significant decrease in latency to the hidden platform across training days, and differences in dwell time between target and non-target quadrants – demonstrating robust spatial learning and memory performance in the MWT (Figure 1B). To examine context fear learning in the CFD paradigm, we quantified dwell time in each context during a pre-exposure and preference test period before and after foot-shock conditioning, respectively (Figure 1C-D). If rats learn to associate fear with the shock-paired context, we should observe no differences in dwell time during pre-exposure, but significant differences during the preference test. Indeed, we found a significant change in dwell time between pre-exposure and preference testing, wherein rats learned to avoid the shock-associated context following foot shock conditioning (Figure 1D). Given that rats expressed robust spatial- and fear-based learning and memory in the MWT and CFD, we then sought to characterize the pattern activity sparsity in the hippocampus during performance of each task.

In both the MWT and CFD, we observed strong differences in activity sparsity across hippocampal subregions and the dorsoventral axis (Figure 2A-B). Namely, we found sparser activity in ventral than dorsal subregions, and that the proportion of active cells increased across the trisynaptic circuit (DG < CA3 < CA1; Figure 2B). Critically, we found no eiect of task on sparsity or interactions with hippocampal subregion or axis. To measure the similarity of sparsity across all hippocampal subregions and the dorsoventral axis the two tasks, we computed a representational dissimilarity matrix (RDM) from the absolute diierence in sparsity of all pairwise comparisons in the MWT and CFD (Figure 2C). We then calculated the rank-order correlation (Kendall’s Tau) of RDMs from the MWT and CFD, which showed that the pattern of sparsity is highly conserved across behavioral tasks (Figure 2D). Finally, to relate our measurement of sparsity to memory storage capacity in a biologically plausible Hebbian synapse, we estimated the Hebb-Marr storage capacity (Marr, 1971; Thome et al., 2017) (*Mmax*; see Detailed Methods) directly from measured levels of sparsity across tasks (Figure 2E). As expected (Marr, 1971; Thome et al., 2017), this analysis revealed that storage capacity is greater for sparser populations, wherein ventral DG has the greatest capacity to store distinct patterns of activity, and dorsal CA1 has the lowest capacity for independent pattern storage.

**Figure 2.**
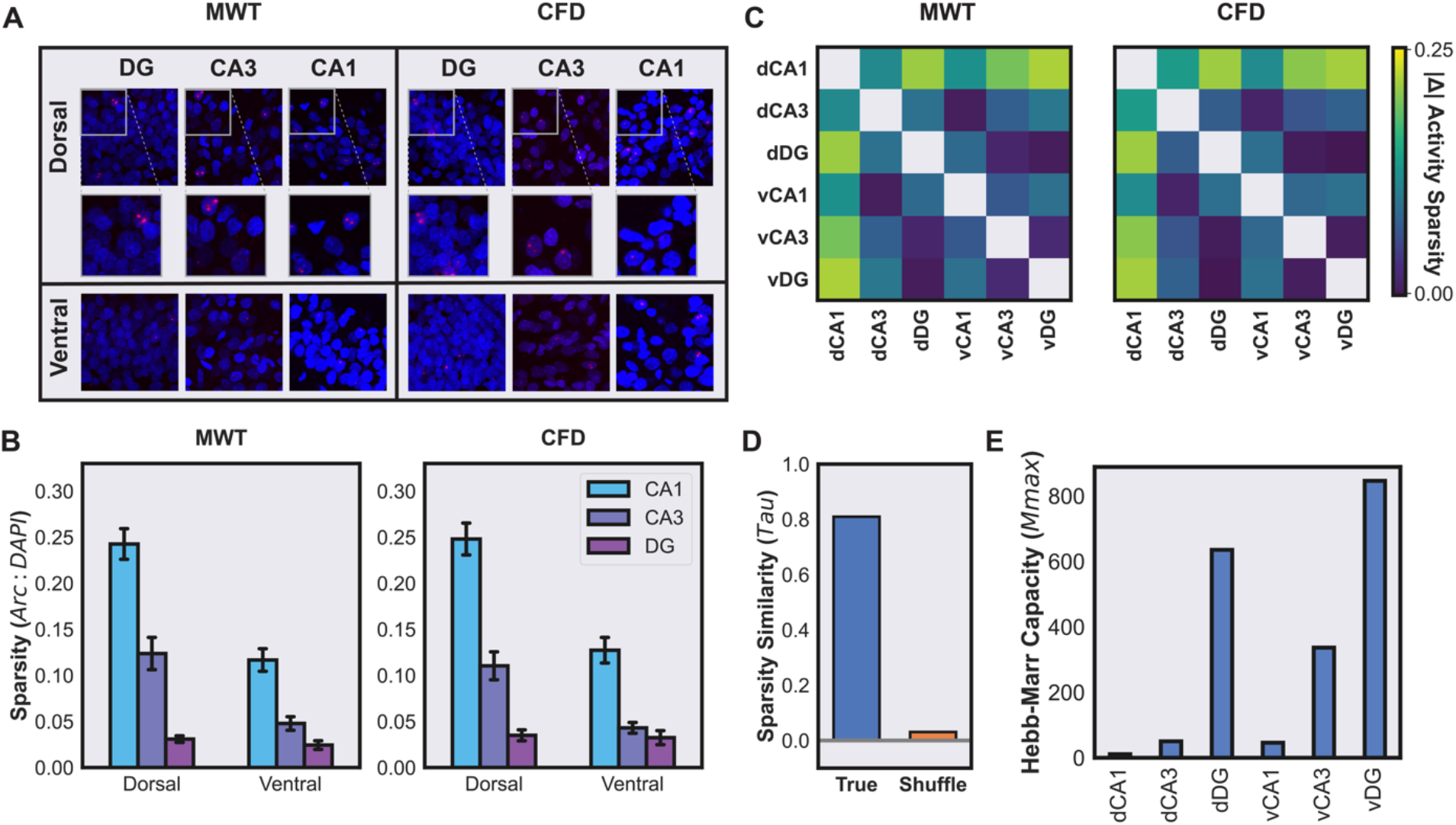
Activity sparsity is task-invariant across the trisynaptic circuit and long axis. **(A)** Representative z-stack image projections of *Arc* (red) and DAPI (blue) labelling from DG (left), CA3 (middle), and CA1 (right) in dorsal (upper) and ventral (lower) hippocampus following performance of the MWT and CFD. The inset for dorsal regions shows a 2x magnification of the selected box. **(B)** A three-way ANOVA revealed a significant effect of subregion (*F*=172.9828, *P*<0.0001), dorsoventral axis (*F*=95.4128, *P*<0.0001), and subregion X axis interaction (*F*=25.1513, *P*<0.0001), but no effect of task (*F*=0.0111, *P*=0.9161), task X subregion (*F*=0.6108, *P*=0.5444), task X axis (*F*=0.1825, *P*=0.6699), or task X subregion X axis interaction (*F=*0.0099, *P=*0.9901). Since we did not find any significant effects of task, we then combined data from both tasks to perform post-hoc comparisons across subregions and the dorsoventral axis. This revealed significant differences in sparsity across the trisynaptic circuit, wherein DG activity is more sparse than CA3 (*t*=-5.7695, *P*<0.0001) and CA1 (*t*=-12.9823, *P*<0.0001), and CA3 activity is more sparse than CA1 (*t*=-10.8354, *P*<0.0001). Comparing the dorsoventral axis of each subregion, we found significant differences in activity sparsity between dorsal and ventral in CA3 (*t*=6.6052, *P*<0.0001) and CA1 (*t*=9.2705, *P*<0.0001), but not DG (*t=*1.2129, *P*<0.0001). **(C)** Representational dissimilarity matrices (RDMs) were calculated from the absolute difference in average activity sparsity of each subregion in the dorsal and ventral hippocampus, wherein the absolute difference between all pairs of regions activity sparsity is shown. Dark colors indicate a similar level of sparsity, and brighter colors indicate greater differences across comparisons. **(D)** To measure the similarity of sparsity across regions in both tasks (MWT and CFD), we computed the rank-order correlation (Kendall’s Tau) the RDMs shown in (C), as well as the average correlation of repeated shuffles of the same RDMs (chance level). The graph illustrates that sparsity is highly correlated across tasks (*Tau*=0.8095; *P*<0.0001) and greater than chance (shuffle average: *Tau*=0.0309, *P*=0.4776). **(E)** To estimate memory capacity from activity sparsity, we computed the Hebb-Marr Capacity (*Mmax*; see Detailed Methods) from sparsity estimates in each subregion and across the dorsoventral hippocampal axis. The results illustrate sparsity is inversely related to memory capacity, wherein sparsely representations can store a greater number of orthogonalized activity patterns.

In the present study, we asked if the organization of sparsity in the hippocampal system depends on performance of a specific memory task. Namely, we examined how sparsity varies across the trisynaptic circuit and hippocampal long axis when animals navigate to a hidden goal location or avoid a shock-associated context. We found that activity sparsity in the hippocampus is highly organized and task-invariant, wherein activity increases across the trisynaptic circuit, and decreases ventrally. To our knowledge, this is the first study to measure sparsity across the entire dorsoventral axis of the trisynaptic circuit during performance of a memory task. Future work should determine the mechanisms that organize sparsity in this canonical circuit, and its consequences for behaviour in both biological and artificial systems.

It is perhaps surprising that we observed a highly similar pattern of sparsity while animals performed spatial- and fear-based memory tasks, given a growing number of studies demonstrating functional differences in these regions for such aspects of cognition (Bannerman et al., 2004; Fanselow & Dong, 2010). However, several studies have shown that dorsal and ventral regions of the hippocampus have an important role in both spatial and fear memory (Ferbinteanu et al., 2003; McDonald et al., 2018; Sutherland et al., 2008), and that lesions to ventral and dorsal hippocampus impact coarse and precise navigation early and in late training (McDonald et al., 2018; Ruediger et al., 2012), respectively. With such evidence, a spatial versus fear-based cognitive division does not adequately describe functional differences across the long axis of the hippocampus.

While we found task-invariant patterns of sparsity across the long axis of the trisynaptic circuit, such regions also have well-described differences in anatomical connectivity to diverse cortical and subcortical structures (Fanselow & Dong, 2010; Moser & Moser, 1998). The variation in sparsity we have observed across the dorsoventral hippocampal axis likely has important consequences for neural coding in down-stream regions. A highly sparse representation, such as that observed in ventral hippocampal subregions, could be rapidly decoded in downstream target areas; for example, to distinguish between threatening and non-threatening contexts. Due to its highly sparse activity, one could expect that the ventral hippocampal representation would be less likely to suffer from catastrophic interference and forgetting (Marr, 1971; Mcclelland et al., 1995). Indeed, it was recently observed that the ventral hippocampus is necessary for discrimination between highly similar contexts (McDonald et al., 2018) and that contexts can be more rapidly decoded from ventral than dorsal hippocampal population activity (Rozeske et al., 2023).

In contrast to our findings that sparsity is task-invariant, prior work has shown that the richness of sensory conditions impacts the sparsity of hippocampal population activity. It has been well-described that animals in a cue-poor, home-cage environment have sparser hippocampal activity compared to those navigating freely or performing a memory task (Chawla et al., 2005; Guzowski et al., 1999; Lee et al., 2019; Lee et al., 2018; Lee et al., 2022; Witharana et al., 2016). In addition, recent work also reported that navigation in virtual environments results in sparser representation than real-world navigation, likely due to attenuated sensory conditions in virtual reality (Thome et al., 2017). We speculate that sparsity is in part determined by the richness of sensory signals, but anatomical constraints set an upper-bound on sparsity. Future work should address how diverse sensory conditions determine activity sparsity across distributed brain areas and its neurobiological constraints.

To disentangle the neurobiological factors controlling activity sparsity across distributed areas, future work should directly manipulate factors hypothesized to control sparsity, such as cellular excitability and afferent connectivity. Recently developed opto- and chemo-genetic tools now allow for the direct manipulation of cellular excitability and targeting of specific projections onto target populations (Rost et al., 2022; Sternson & Roth, 2014). In combination with tasks to probe canonical computations such as pattern separation (Cayco-Gajic & Silver, 2019), pattern completion, and task transfer, it will be important directly examine how sparsity causally contributes to the ability of animals to perform tasks that probe such cognitive functions.

Importantly, behavioral factors, such as environmental enrichment, could also impact sparsity of neural coding in distributed systems (Saxena & McNaughton, 2024). We anticipate that sparsity is titrated to contexts in which animals commonly navigate, learn and remember – daily exposure to a cue-rich rearing environment could result in sparser representation during cognitive task performance than a cue-poor environment (Bilkey et al., 2017), possibly through recruitment of inhibition along with plasticity mechanisms. Advancing our understanding of the biological constraints for activity sparsity could also inform brain-inspired models of sparse activation functions that could have important implications for developments in machine learning and artificial neural networks.

Indeed, sparsity is an active, growing area of study in machine learning and artificial intelligence in supervised, self-supervised and reinforcement learning domains (Liu et al., 2019). For example, recent studies in reinforcement learning and supervised learning in artificial neural networks has revealed that specific, sparse activation functions can aid in continual learning (Wang et al., 2024). Interestingly, activation functions that produce a one-to-many connectivity result in more robust representation learning and knowledge transfer across tasks (Wang et al., 2024). In addition, some work in the supervised setting has also shown that gradients of sparsity emerge naturally with specific architectures and learning rules (Li et al., 2023). It will be valuable for future work to explore biologically plausible learning mechanisms that produce organized patterns of sparsity that are task-invariant, such as the patterns we have observed across the trisynaptic circuit and dorsoventral axis of the hippocampus, and their consequences for intelligent behavior in artificial systems.

## Detailed Methods

The animals and behavioural data in the present study were described previously Lee et al. (2022), along with all behavioral procedures, measurements, and tissue processing methods.

### Animals

The University of Lethbridge Animal Welfare Committee approved all procedures used in the present experiments, which also meet the Canadian Council of Animal Care guidelines. A total of 24 Long Evans rats weighing between 300 and 350 g were used in the present experiments, including 12 females and 12 males (Charles River, Raleigh, NC). Previous work has shown such group sizes provide adequate statistical power for within-animal and across-group comparisons for behavioural and cellular levels of analysis. Following their arrival at the University of Lethbridge, animals were allowed at least 1 week to acclimatize to colony room conditions and were handled 5 min each day by the experimenter for 5 days before the start of the experiment.

### Experimental Design

To match behavioural experience across animals, male and female rats were equally divided into cohorts and trained in the MWT and CFD task in counterbalanced order before final testing and perfusion (Fig. 1). Our rationale for training animals both tasks was two-fold: 1) to ensure similar cumulative experience prior to final testing and perfusion that might otherwise aiect *Arc* expression; 2) to ensure similar performance of groups in MWT and CFD performance. Each animal experienced the MWT and CFD task for 4 days each, with one day of rest between paradigms. During training in the MWT, rats were transported from the animal colony to the behavioural testing room in covered cages on a cart. Surrounding the swimming pool were several posters on the walls, a table, and computer rack that served as allocentric cues to help animals navigate. During each session, animals were monitored, and relevant behavioural variables (latency and quadrant dwelling) were calculated using EthoVision XT software (Noldus) from an overhead behavioural camera. The MWT apparatus consisted of a 2-meter diameter pool filled with room temperature water (∼25 C) that was made opaque using white non-toxic tempura paint. On days 1–3, rats were given 8 trials (maximum 60 s) starting randomly from one of the four cardinal positions at the edge of the pool to locate a hidden platform approximately 5 cm beneath the water surface located in the center of the Northwest quadrant. If the animal did not locate the hidden platform within 60 s they were placed onto the platform by the experimenter. Animals were then allowed 10 s to remain on the platform before placement back into their holding cage by the experimenter for an approximate 5-minute intertrial interval. Following completion of the 8 swim trials each day, rats were returned to their colony room for approximately 24 h prior to subsequent behavioural training or testing. During the final testing day in the MWT, animals were individually transported in covered cages and given 4 swim trials with a 2-minute intertrial interval for a total 10-minute testing period. Half of the animals from female and male groups were euthanized and perfused following MWT testing.

In the CFD task, rats were individually transported in a covered holding cage by the experimenter to a room with several posters on the walls, a storage shelf, and the CFD apparatus, which consisted of two conditioning chambers (contexts) and connecting alleyway. One context was a black triangle that was 61 cm long, 61 cm wide, and 30 cm high with stainless steel rod flooring, and was scented with banana (Isoamyl Acetate, Sigma) located in a perforated pill bottle at the top right corner of the chamber. The other context was a white square that was 41 cm long, 41 cm wide, and 30 cm high with stainless steel rod flooring and was scented with Eucalyptus (Vic’s VapoRub©) located in a perforated pill bottle inserted through the top left corner of the chamber. During each session, animal behaviour was recorded from a tripod-mounted camera from underneath the apparatus through a transparent table. During pre-exposure on day 1 of the CFD task, animals were introduced to the apparatus through the connecting alleyway and allowed to freely explore both contexts for 10 min. On subsequent days 2 and 3, animals were conditioned in shock-paired and no-shock contexts in counterbalanced order. During no-shock conditioning, animals were placed in their no-shock context for 5 min and allowed to explore freely. For shock-paired conditioning, animals were transported to a distinct and separate room containing the same apparatus and placed into their shock-paired context. Shock conditioning was performed in a distinct room with the same apparatus to promote local-context based fear, rather than room-based, spatial fear learning, based on previous studies that have demonstrated room transfer drives remapping in hippocampal place cells. During shock conditioning, the stainless-steel rod flooring was connected to a Lafayette Instrument Stimtek SGCG1 through a custom shock harness, and 2-second, 1.0 mA scrambled foot shocks were delivered at the 2nd, 3rd, and 4th minute. After an additional 58 s, animals were removed from the shock-paired context and returned to their home cage. On day 4 of the CFD task, animals were returned to the original training room, and introduced to the apparatus through the connecting alleyway and allowed 10 min to explore both contexts. Half of female and male groups were sacrificed and perfused following final preference CFD testing (Fig. 1E). Dwell time in each context was calculated by a trained observer blind to conditions of sex and testing order from video data of the pre-exposure and preference testing epochs and was defined as the presence of both forepaws in a context. Following final testing in either the MWT or CFD, animals were returned to their holding cage for 1 min, and then given an overdose intraperitoneal injection of sodium pentobarbital. They were then perfused 5–10 min after the completion of behavioural testing in either task, and then decapitated and had their brains extracted for subsequent tissue processing. This timeline was chosen based on previous studies demonstrating that behaviour-driven *Arc* expression is maximal 5–10 min after a learning or remembering episode.

### Tissue Processing

The methods used for Arc visualization with FISH are identical to those in previous studies from our group on *Arc* expression in the MWT. Following fixation and sectioning at 50 μm thickness in a 12-section series using a freezing-sliding microtome, samples were stored at -80 C until fluorescent *in situ* hybridization (FISH) tissue processing. *Arc* riboprobes were designed to detect intronic mRNA sequences, and thus nuclear rather than cytoplasmic expression. Primers flanking *Arc* intron 1, exon 2, and intron 2 were designed using online software (National Center for Biotechnology Information Primer-Blast; credit to A. M. Demchuk, University of Lethbridge). The exact sequences of the primers and base pair designations follow those of the GenBank accession number NC_005106: 5’-CTTAGAGTTGGGGGAGGGCAGCAG-3’ (forward primer, base pairs 2022 - 2045) and 5’-ATTAACCCTCACTAAAGGGCCCTGGGGCCTGTCAGATAGCC-3’ (reverse primer tagged with T3 polymerase binding site on 5’ end, base pairs 2445–2466). The polymerase chain reaction (PCR) was performed on a genomic rat DNA template using a Taq PCR Kit (New England Biolabs), and the subsequent PCR product was purified using a Qiagen PCR Purification Kit (Life Technologies, Inc.). The MAXIscript T3 transcription kit (Life Technologies, Inc.) and DIG RNA Labeling Mix (Roche Diagnostics) were used to generate DIG-labelled Arc intron-specific antisense riboprobes from PCR templates. Riboprobes were then purified with mini QuickSpin columns (Roche Diagnostics), and FISH was performed on slide-mounted tissue as described previously. Briefly, DIG-labelled Arc riboprobe signal was amplified with anti-DIG-POD (1:300; Roche Diagnostics), Tyramide Signal Amplification (TSA) Biotin Tyramide Reagent Pack (1:100; PerkinElmer), and Streptavidin-Texas Red (1:200; PerkinElmer). Cell nuclei were then counterstained with DAPI (1:2000; Sigma-Aldrich).

### Functional molecular imaging

The approach used for quantification of Arc and DAPI labels was identical to the methods described in Lee et al. (2019)(Lee et al., 2019) and Lee et al. (2022)(Lee et al., 2022). Briefly, observers blind to experimental conditions of each animal quantified DAPI and Arc expression using the optical fractionator method in StereoInvestigator software (version 10.54, MBF Bioscience, VT) from confocal z-stack images collected on an Olympus FV1000 microscope equipped with Fluoview software (version 4.0, Olympus, Shinjuku, Japan). Unilateral traces of DG, CA3, and CA1 were created at 20X magnification on each section, and counting frames were automatically positioned according systematic-random sampling procedures with a 150 ×150 μm grid over the traces for each subregion. A series of seven z-stack images at 512 ×512-pixel resolution was collected at each sampling site with a 60X oil-immersion objective starting at the top of the section every 2 μm for a total 14 μm sampling distance in the z-plane. Image thresholds were set at 700 HV ±20 and 550 HV ±20, respectively in DAPI and Texas Red channels, and kept constant across imaging each section series such that small Arc foci (2–3 pixels in diameter) and DAPI labels could be clearly identified. Digital z-stack images were then imported into StereoInvestigator software, such that the top image from each stack fell above and the final image below a 10-μm height of the optical dissector volume. *Arc* and DAPI were then counted according to optical fractionator inclusion–exclusion criteria at each cell’s widest point in a 30 ×30 × 10 μm fractionator probe. One animal was excluded from *Arc* quantification due to tissue damage from experimenter error.

### Statistics

All statistical analyses were performed in Python 3.7 using Numpy (v1.24.4), Pandas (v2.0.3), and Statsmodels (v0.14.1) libraries. Data visualization was performed with the libraries Matplotlib (version 3.7.5) and Seaborn (version 0.13.2) in Python. All data and code to perform all analyses and to reproduce the main figures can be found at: https://github.com/jquinnlee/HPCArc. To generate dissimilarity matrices for sparsity comparison across regions and tasks, we measured the absolute diierence of average sparsity in each subregion for MWT and CFD cohorts for all pairwise comparisons. We then calculated the rank-order correlation (Kendall’s Tau) of dissimilarity matrices for both tasks (True) and performed the same comparison for 1e3 random permutations of the same matrices (Shuffle). Storage capacity for a Hebb-Marr memory was estimated from sparsity measurements as described previously in Thome et al. (2017)(Thome et al., 2017), where:

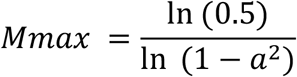

## Acknowledgements

We thank Aubrey Demchuk, Valérie Lapointe, and Maurice Needham for their technical assistance with this work. This work was supported by the Natural Sciences and Engineering Research Council of Canada grant awarded to Robert J. McDonald and the Chinook Summer Research Studentship awarded to Matt Nielsen.

